# Methylation variability and LINE-1 activation in multiple myeloma

**DOI:** 10.1101/2025.07.22.666173

**Authors:** Qianhui Wan, Amy Leung, Mahek Vinod Bhandari, Hiroyuki Kato, Joo Y. Song, Dustin E. Schones

**Author notes:** **Corresponding Author:** Dustin E. Schones, Ph.D. Department of Diabetes Complications and Metabolism City of Hope Duarte, CA, 91010.

## Abstract

Multiple myeloma (MM) is a type of hematological cancer that arises from uncontrolled proliferation of plasma cells. In addition to frequent genetic mutations, malignant plasma cells are characterized by alterations to the epigenome. Myeloma cells display a genome-wide loss of DNA methylation and a corresponding increase in ‘active’ chromatin modifications. The dramatic epigenetic remodeling that occurs in cancer genomes is associated with loss of silencing at transposable elements, which can impact genome regulation. Through paired epigenome and transcriptome profiling of patient derived MM samples, we have found that loss of DNA methylation in MM genomes results in the formation of partially methylated domains that are variable across patients. This loss of DNA methylation coincides with the expression of hundreds of transcripts driven by LINE-1 (L1) retrotransposons that are epigenetically silenced in normal cells. MM samples can be stratified based on L1 activity with distinct gene expression signatures. The high L1 samples are characterized by a more proliferative, less differentiated state as well as inhibition of interferon and genome defense pathways. Several L1 promoters generate chimeric transcripts with adjacent oncogenes. We further find that KRAB-zinc finger proteins (KZFPs) that are responsible for the epigenetic silencing of L1s have abnormally low abundance in MM samples with high L1 activity. These results indicate that cell proliferation in MM is associated with a loss of KZFP expression and activation of L1 elements.

**Significance:** Epigenome and transcriptional profiling of patient-derived multiple myeloma samples reveals heterogeneous loss of DNA methylation, reduction of KZFP expression and LINE-1 activation as a clinical indicator of progression.

## Introduction

Multiple myeloma (MM) is a hematologic malignancy that accounts for approximately 15% of all blood-related cancers (1). It arises from malignant transformation and uncontrolled clonal expansion of plasma cells, a subset of differentiated B-lymphocytes responsible for immunoglobulin production. The infiltration of these malignant plasma cells into the bone marrow leads to various clinical manifestations, including bone lesions, anemia, renal impairment, and immune dysregulation (2). The disease is characterized by complex genetic and epigenetic alterations that contribute to heterogeneity, pathogenesis, and therapeutic resistance.

Plasmablasts represent an immature and highly proliferative stage of B cell differentiation, serving as precursors to mature plasma cells. Under normal physiological conditions, plasmablasts undergo apoptosis after a few days or differentiate into long-lived plasma cells that continue antibody secretion (3). However, in certain pathological contexts, the dysregulation of plasmablast differentiation and survival can contribute to malignancies. Plasmablastic multiple myeloma (PMM) was first defined in 1985 as a subtype of MM characterized by an enrichment of plasmablasts, comprising more than 2% of bone marrow cellularity (4). Compared to conventional MM, PMM exhibits distinct biological and clinical features, including an aggressive disease course, elevated proliferative capacity, and poorer overall prognosis (5). The aggressive nature of PMM presents significant challenges in both diagnosis and treatment, necessitating further investigation into its underlying molecular mechanisms.

As is common in many types of cancer, epigenetic alterations are common in B cell malignancies, with MM having the most dramatic loss of methylation of B cell tumor types (6). MM cells generally have epigenetic profiles similar to undifferentiated cells with malignant plasma cells with DNA methylation alterations impacting the utilization of regulatory regions (6). One of the primary roles of DNA methylation in mammalian genomes is in the epigenetic repression of transposable elements (TEs) (7). More than half of the human genome consists of repetitive sequences derived from TEs, the vast majority of these being retrotransposons (8). These retroelements primarily consist of long-terminal repeat (LTR) containing endogenous retroviruses (ERVs) as well as short and long interspersed nuclear elements (SINEs and LINEs). While much of the genetic content of these elements has eroded throughout evolution, a substantial number have retained the capability of impacting genome regulation. TEs are targeted by KRAB-containing zinc finger proteins (KZFPs) for epigenetic repression in development (9). KZFP genes themselves have evolved alongside repeat elements to maintain repression of TEs. The general epigenome dysfunction that is common in cancer cells can lead to loss of suppression of TEs. This can impact the host genome in several ways. Loss of silencing of LTR elements in cancer can lead to repurposing of regulatory elements in cancer, a process known as “onco-exaptation” (10). Similarly, loss of repression at LINE-1 elements can lead to active transposition events, which are common in cancers (11).

Several key unknowns remain in the context of multiple myeloma (MM) and its aggressive variant, plasmablastic multiple myeloma (PMM). First, MM itself is a highly heterogeneous disease at both the genetic and transcriptomic levels, yet the mechanisms driving this heterogeneity are still not fully understood (12). The more aggressive PMM variant is significantly less studied than MM overall, limiting our ability to draw parallels or identify distinguishing molecular features (13). The DNA methylation landscape of PMM has not been systematically characterized, leaving its epigenetic regulation poorly defined. The expression patterns of KRAB zinc finger proteins (KZFPs) are also largely unstudied in PMM. Lastly, the behavior and regulation of TEs in MM remains largely unexplored, in contrast to what is known for other cancers (12).

We report here a comprehensive investigation into the DNA methylation landscape and TE expression patterns in PMM and a comparative analysis with MM. Using a novel approach specifically developed to analyze transcriptome data from formalin-fixed paraffin-embedded (FFPE) tissues, we identified extensive activation of LINE-1 (L1) elements in PMM samples. Further analysis of 764 MM transcriptome profiles from the MMRF-CoMMpass study revealed similar trends. By elucidating the epigenetic mechanisms underlying PMM, our findings provide novel insights into the regulatory architecture of the disease and identify potential therapeutic targets for this aggressive myeloma subtype.

## Materials and Methods

### Tissue collection and processing

Formalin-fixed, paraffin-embedded (FFPE) tissue samples from plasmablastic multiple myeloma (PMM) patients (N = 15) were collected following Institutional Review Board approval (IRB #18136) and selected based on confirmed diagnosis. Cases defined as plasmablastic myeloma showed >20% large, atypical plasma cells with prominent nucleolus (plasmablasts). These cases were selected from archival material in the Department of Pathology at City of Hope. Total RNA was extracted using the Qiagen RNeasy FFPE Kit, assessed for integrity (RIN > 10), and sequenced using paired-end (150 bp), dUTP-based, strand-specific (reverse-stranded) RNA sequencing on an Illumina NovaSeq 6000 platform, targeting 200 million reads per sample. Matched genomic DNA was extracted using the Qiagen QIAamp DNA FFPE Tissue Kit, bisulfite-converted (>99% efficiency using the Zymo EZ DNA Methylation-Gold Kit), and sequenced via whole-genome bisulfite sequencing (WGBS) on an Illumina NovaSeq 6000.

### Preprocessing of RNA sequencing data

For alignment of fastq files to hg38 reference genome (chromosome 1 to 22, X, Y and M considered), 2-pass alignment with *STAR* (14) was applied with the parameters: “--twopassMode Basic --outFilterMultimapNmax 20 --alignSJoverhangMin 8 -- alignSJDBoverhangMin 1 --outFilterMismatchNmax 999 –outFilterMismatchNoverLmax 0.1 --alignIntronMin 20 --alignIntronMax 1000000 --alignMatesGapMax 1000000 -- outFilterType BySJout --outFilterScoreMinOverLread 0.33 -- outFilterMatchNminOverLread 0.33 --limitSjdbInsertNsj 1200000 --outSAMstrandField intronMotif --outFilterIntronMotifs None --alignSoftClipAtReferenceEnds Yes -- quantMode TranscriptomeSAM GeneCounts --outSAMtype BAM Unsorted -- outSAMunmapped Within --genomeLoad NoSharedMemory --chimSegmentMin 15 -- chimJunctionOverhangMin 15 --chimOutType Junctions SeparateSAMold WithinBAM SoftClip --chimOutJunctionFormat 1 --chimMainSegmentMultNmax 1”. *Picard* was used to identify and remove duplicate reads (15). For stranded RNA-seq, BAM files for paired reads were generated separately for the Crick and Watson strands. For non-stranded RNA-seq (from MMRF), a single BAM file was generated for each sample. Uniquely mapped reads were filtered using *featureCounts* package (16) and only uniquely mapped reads marked with NH:i:1 were used for the activation score described below.

### Preprocessing of WGBS data

The *trim_galore* package (17) was used to trim adapters from each FASTQ file with parameters: “--paired --clip_R1 8 --clip_R2 8 --three_prime_clip_R1 9 -- three_prime_clip_R2 9”. Trimmed FASTQ files were aligned to the hg38 genome using the *Bismark* package (18) with parameters: “--bowtie1 --non_bs_mm --unmapped -- phred33-quals”, and duplicate reads were identified and removed using *Picard* (15). The coverage between samples was normalized by *methylKit* R package (19). This method corrects for differences in sequencing depth across samples by computing a scaling factor for each sample based on the median of the coverage values. Each sample’s coverage is then scaled to bring all samples to the same overall coverage distribution. The CpG methylation percentage was calculated as the number of unconverted cytosines (Cs) divided by the total number of reads covering those sites. DNA methylation sites with coverage of less than 5 reads were filtered out using the *methylKit* R package (19).

### Annotation of transcripts initiated from TE regions

Human hg38 genome transcript reference was obtained from GENCODE database (https://www.gencodegenes.org/human/release_46.html). Annotations of hg38 genome TE regions from RepeatMasker (20) were obtained from the UCSC Genome Browser (https://genome.ucsc.edu/cgi-bin/hgTables). Transcription start sites (TSSs) were extracted according to the corresponding strand information from GENCODE transcript annotation. The transcripts with TSSs located in TE regions were selected as annotated known TE transcripts. Full length L1 regions on hg38 reference genome were obtained from L1baseV2 (21) (http://l1base.charite.de/l1base.php). LINE1 elements longer than 5900 bp from RepeatMasker were selected for downstream statistical analysis according to L1 length distribution.

### Differential gene expression analysis

The *featureCounts* package (16) was first used to quantify fragments for each gene from BAM files, excluding low-quality reads, multi-mapped reads, and ambiguous reads. After generating the count matrix for all samples using *featureCounts*, we constructed a list containing both the sample metadata and the count matrix using the *edgeR* package (22). Genes were retained if they had a log counts per million (CPM) > 1 in at least as many samples as the size of the smallest experimental group, for example, genes with CPM > 1 in at least 3 samples were retained for PMM samples, and those with CPM > 1 in at least 149 samples were retained for MM samples. A multivariate linear model was then applied to voom-transformed, filtered raw counts to identify differentially expressed genes between groups using the *limma* package (23). Differentially expressed genes were defined as absolute logFC >1 and false discovery rate (FDR) <0.05. A filtered gene count matrix, normalized using the trimmed mean of M values (TMM) method (24) to adjust for library size differences between samples, was used for visualization (24).

### Definition and calculation of activation score

Previous efforts identifying transposon-derived transcripts have mostly utilized transcriptome data generated from frozen cancer tissues. With short (50-100bp) reads in these datasets, traditional transcript assembly methods such as *StringTie* (25) and *Salmon* (26) with reference guided parameter settings have been shown to identify transcripts that arise from transposon promoters (27). However, these methods are not optimal for FFPE samples where extracted RNA is typically fragmented and assembled transcripts are incomplete. We therefore developed a novel pipeline to distinguish active transposon promoters in FFPE samples. We generated a transposon promoter identifier based upon expected read alignment profiles of transcription originating from a transposon promoter vs intronic “read-through” transcription. We reasoned that if a transposon contains an active promoter, the number of reads aligning downstream will be very high compared to the number of reads aligning upstream of the transposon – the ratio between those two numbers (downstream reads/upstream reads) is >>1. Alternatively, if a transposon contains reads because it sits within an independently transcribed unit (e.g. an intron), the ratio would be ∼1. To choose a cut-off for the ratio that accurately reflects the sample data, we generated expected ratios of random intronic regions (n=5683690). From this, we used the top 1% of the ratios from random regions as the threshold to call RTEs as promoters.

### Weighted gene co-expression network analysis (WGCNA)

To investigate genes correlated with KZFP expression, we performed WGCNA (28) on PMM and bone marros plasma cell (BMPC) samples (n = 11). Briefly, WGCNA calculates pairwise correlations between gene expression profiles (n = 26,925 genes) and constructs a weighted adjacency matrix. A soft-thresholding power of 12 was selected to approximate scale-free topology. The topological overlap matrix (TOM) was computed to measure network interconnectedness, and hierarchical clustering was applied to group genes into modules based on their co-expression patterns. Gene modules were defined using the dynamic tree cut method, and modules with similar expression patterns (eigengene correlation ≥ 0.60) were merged using a cut height of 0.40 resulting in 19 modules for further analysis. The co-expression network involving ZNF141 and ZNF382 within the lightskyblue module was filtered out and then gene pairs with the top 25% of TOM similarity values (edge thickness) were retained to emphasize strong network connectivity. The resulting network was visualized using *Cytoscape* software (29) to illustrate interactions within the module.

### Identification of partially methylated domains (PMDs)

The identification of PMDs was performed using a two-state Hidden Markov Model (HMM) implemented in the *dnmtools* (30) rather than at the single CpG level. Given that PMDs are large, megabase-scale regions characterized by widespread methylation loss and high methylation variability at individual CpG sites, each state in the HMM reflects the weighted average methylation level of a genomic region rather than discrete CpG sites. For PMM samples, we additionally derived a consensus PMD state by integrating data across all PMM samples, facilitating a unified definition of PMD regions.

### GSEA analysis

Hallmark gene sets for GSEA analysis were retrieved from the GSEA database using the *msigdbr* R package(31). GSEA analysis was then performed using the *clusterProfiler* R package (32) based on a gene list ranked by the signed significance of gene expression changes between groups. An adjusted *p*-value or FDR < 0.05 was considered statistically significant, and the differential signaling pathways were then visualized with *clusterProfiler* R package.

### Enrichment analysis of repeat subfamily expression

Fisher’s exact test was used to assess the enrichment of each retrotransposon repeat family exhibiting TE activation (activation score > 0.99 quantile of that from random regions) compared to other repeats. FDR correction was applied to adjust *p*-values for multiple comparisons. Enrichment was defined as an observed-to-expected ratio greater than 1 (Observed/Expected > 1) with FDR < 0.05. Observed proportions (numerator of the enrichment score) were calculated as the number of activated TEs in each repeat family divided by the total number of activated TEs. Expected proportions (denominator of the enrichment score) were calculated as the proportion of TEs in each repeat family relative to the total number of TEs.

### Statistical analysis

Statistical significance was assessed using the Wilcoxon rank-sum test for pairwise comparisons unless otherwise indicated. For multi-group comparisons, Kruskal–Wallis or ANOVA tests were applied as appropriate. All *p*-values were adjusted for multiple testing using the Benjamini–Hochberg procedure, and adjusted *p*-values < 0.05 were considered statistically significant.

## Code availability

All analysis scripts and custom pipelines used in this study are publicly available at the following GitHub repository: https://github.com/QianhuiWan/coh_PMM_paper.

## Data availability

RNA-Seq and whole-genome bisulfite sequencing (WGBS) data for PMM samples generated in this study are publicly available in the Gene Expression Omnibus (GEO) under accession number GSE302851, and the raw sequencing data are also available through the Sequence Read Archive (SRA) under BioProject ID: PRJNA1285902.

## Results

### DNA methylation decrease in PMM is associated with increased L1 activity

To characterize epigenetic alterations in PMM, we performed whole-genome bisulfite sequencing (WGBS) from 15 patient-derived samples. The DNA methylation landscape in PMM exhibited a general variable loss of methylation and the formation of partially methylated domains (PMDs) compared to resting B cells (**Fig. 1A**). To verify that the detected PMDs in PMM were in late-replicating genomic regions, the predicted compartment B regions were verified by wave score (late replication represented by low wave score) in HepG2 cells (**Supplementary Fig. S1A**). In PMDs, CpG islands (CGIs) in non-repetitive regions of the genome displayed a general increase in methylation in PMM compared to resting B cells (**Fig. 1B**), consistent with previous reports (33). In contrast, CGIs within transposons have a general loss of methylation in PMM compared to resting B cells (**Fig. 1C**). The same general trends were observed in CGIs but not in PMDs (**Supplementary Fig S1B, C**).

**Fig. 1.**
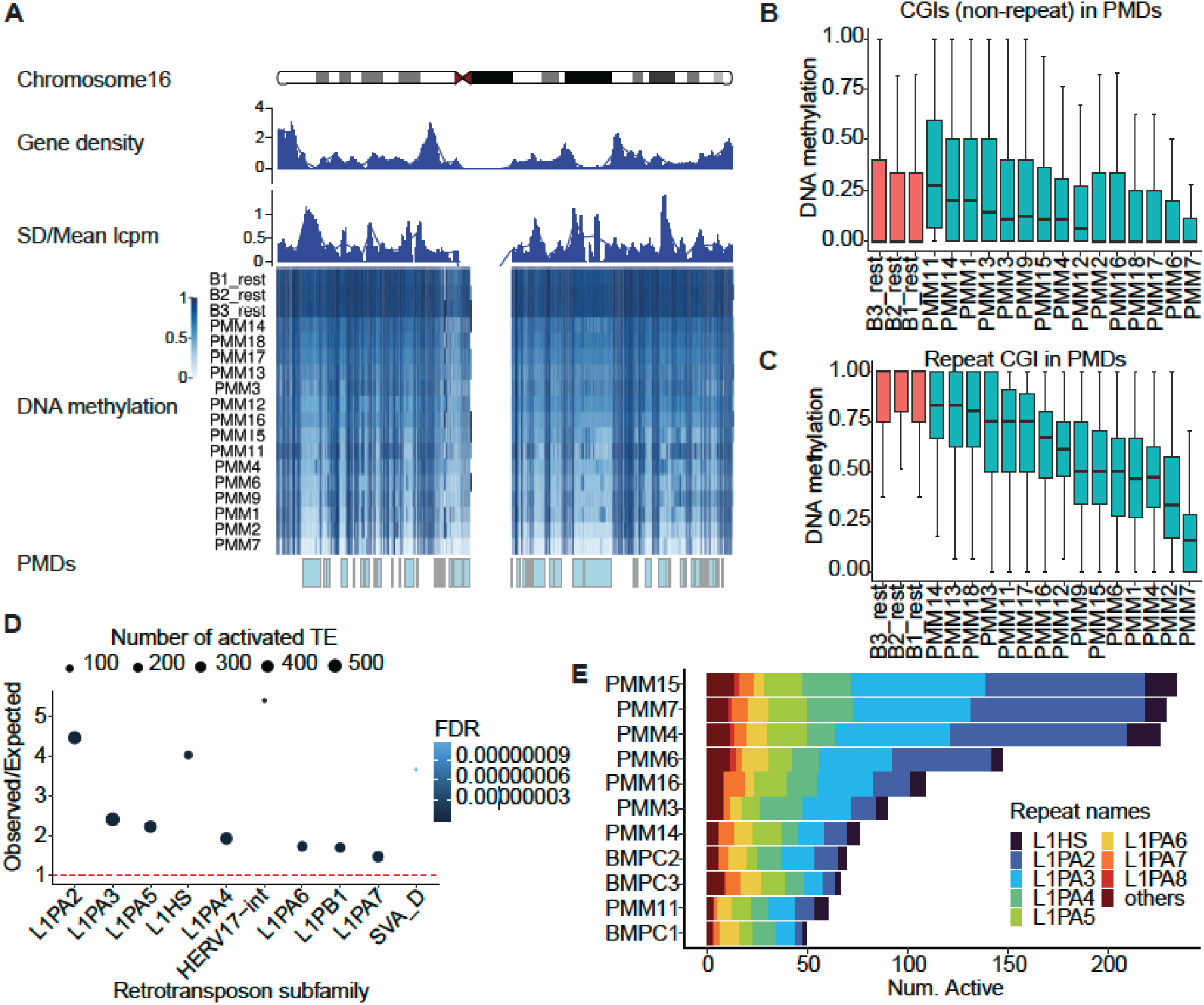
DNA methylation is variably lost at repetitive loci in PMM. **A**, Shown are gene density, standard deviation/mean of lcpm (log2 counts per million), DNA methylation, and PMD regions for Chromosome 16 (as example) across resting B-cells and PMM samples. **B**, Boxplots of DNA methylation percentage at CGIs (non-repeat) in PMDs in each B-cell and PMM sample **C**, Boxplots of DNA methylation percentage at Repeat CGIs in PMDs in resting B-cells and PMM samples. **D**, The top 10 significantly enriched activated repeat subfamilies in PMDs (FDR<0.05 and enrichment >1) ranked by FDR. Shown are observed/expected ratios (enrichment) along with number of activated elements per subfamily. **E**, Number of active L1 elements in each of the samples stratified by repeat subfamily in PMM samples and bone marrow plasma cells (BMPCs).

To assess the impact of DNA methylation loss at repeat elements, we performed RNA-seq for eight patient derived samples and examined the transcriptional activation of TEs (see Materials and Methods for details). Evolutionarily young LINE-1 elements were among the most enriched repeat families activated, followed by HERV, and SVA repeat families (**Fig. 1D**). Among all L1 subfamilies, L1PA2 elements exhibited the greatest degree of activation across different PMM samples (**Fig. 1E**). When examining repeat CpG island DNA methylation variation across the genome, the L1 repeat family remained enriched in variable methylation (coefficient of variation > 0.2 and FDR < 0.05) (**Supplementary Fig. S1D**) and the DNA methylation changes at active L1 CpG islands showed heterogenous loss of methylation (**Supplementary Fig. S1E**).

### Heterogeneous L1 activation in PMM occurs in both sense and antisense strands

L1 activation in PMM was heterogeneous across individual samples, with transcription occurring in both sense and antisense from the L1 promoter CGI (**Fig. 2A**). The distribution of activation scores varied across PMM samples, with a subset showing markedly elevated levels. Genome-wide quantification of the stranded RNA-seq data revealed that while many L1 elements were transcribed in PMM samples in sense from the L1 promoter (n=117; **Fig. 2B, D**), there were markedly more antisense transcripts (n=619; **Fig. 2C, D**). Only a small subset of TE loci (n=10) exhibited simultaneous sense and antisense transcription (**Supplementary Fig. S2A, B**). Benchmarking of two different methods for TE transcript detection showed that the activation score developed in this study achieved a higher F1 score compared to TEprof3 (**Supplementary Fig. S2C**). A cutoff of 5900bp is used to filter for all full length L1 elements based on the length distribution of RepeatMasker annotated L1s (**Supplementary Fig. S2D**).

**Fig. 2.**
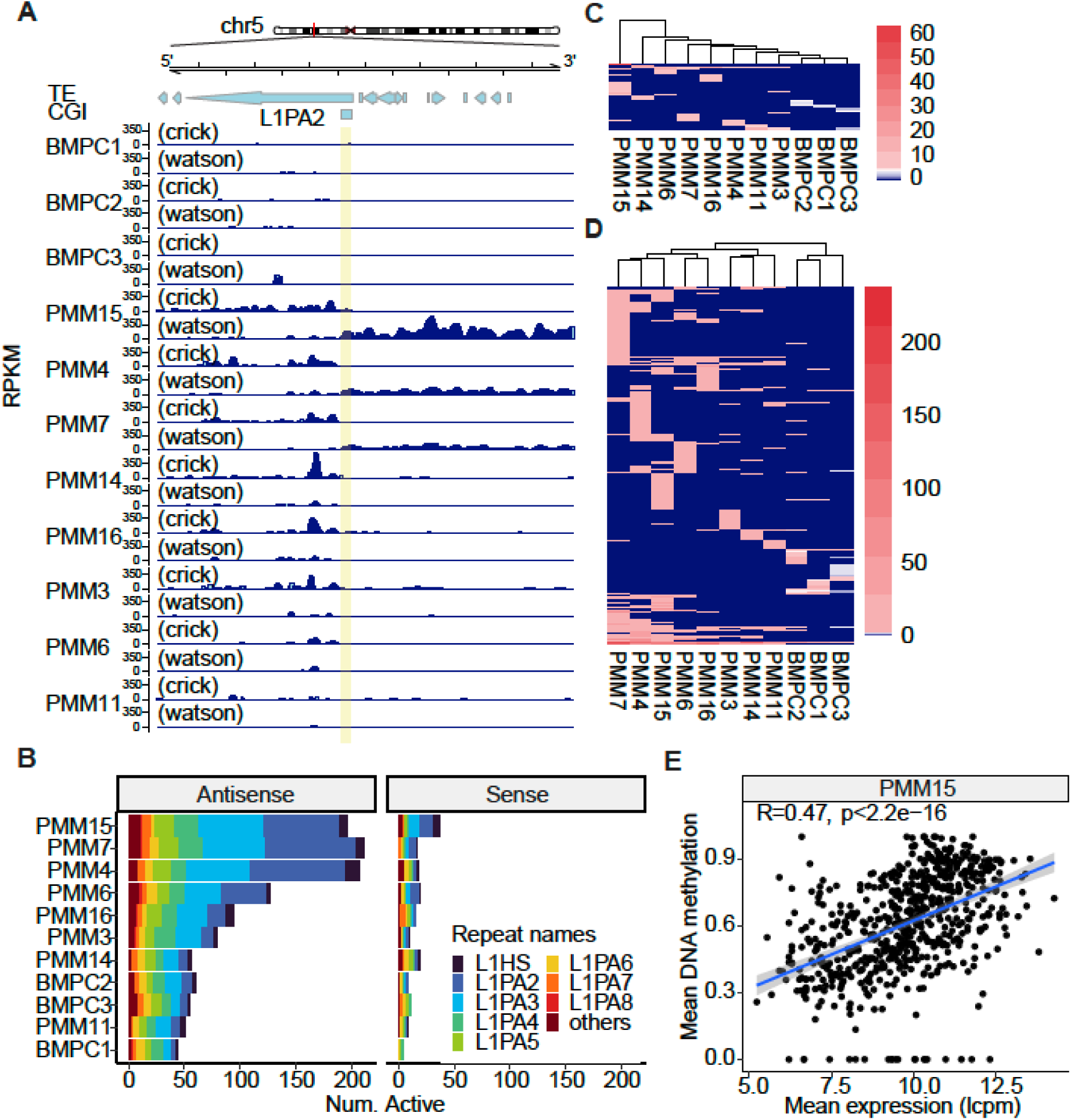
Heterogeneous activation of L1s in PMM. **A,** Stranded RNA-seq signal in BMPCs and PMM samples across a locus on chromosome 5 with TEs annotated (L1PA2 highlighted) and CGI annotated. **B,** Bar graph of number of active L1 elements stratified by subfamily in the antisense (left) and sense orientation (right). (**C, D**), Heatmaps showing BMPCs and PMM samples clustered by activation score for sense (**C**) and antisense (**D**) oriented L1s. **E**, Mean DNA methylation within 10 kb of activated antisense L1 elements in PMM15 sample reveal increased DNA methylation corresponding with increased expression at these regions.

Given the observations of de novo DNA methylation at transcribed regions (34) we wanted to test if activation of L1s would influence DNA methylation in surrounding regions. Average DNA methylation in the 10kb upstream flanking regions of activated antisense L1s in PMM15 sample revealed that more antisense L1 expression correlated with higher DNA methylation (**Fig. 2E**). The same positive correlations were also observed in other PMM samples (**Supplementary Fig. S2E**). The analysis of MEIs in PMM showed that SVA-family MEIs increase in association with L1 activation in PMM (**Supplementary Fig. S2F**).

### LINE1 activation in MM is associated with poor progression

To test the generality of our observation regarding L1 activity in multiple myeloma, we utilized transcriptome profiles from 764 patients that were generated as part of the Multiple Myeloma Research Foundation (MMRF) CoMMpass study (35). Analysis of LINE1 activation in MM revealed substantial interpatient variability. MM patients were stratified into four quantiles (Q1–Q4) based on L1 activation levels, with Q4 patients exhibiting the highest number of expressed L1 elements (**Supplementary Fig. S3A**). Comparison of survival outcomes across these groups showed that higher L1 activity was associated with a lower survival probability (**Fig. 3A**). Among the top 5% of MM patients ranked by active LINE1 elements, ∼60% (22/36) belonged to the PR molecular subtype (n = 51), a highly proliferative subtype with poor survival rates (35) (**Fig. 3B**). Differential gene expression analysis of high L1 vs low L1 samples revealed that MM samples with high L1 activation showed increased E2F pathway activity as well as inhibition of IFN signaling and p53-related processes (**Fig. 3C**). Gene set enrichment analysis of genes ranked by the significance of expression differences between the PMM and BMPC sample groups identified upregulation of epithelial-mesenchymal transition (EMT), and E2F targets pathways in PMM compared to BMPCs. Similar to the MM samples (Q4 vs. Q1), IFN-γ and p53-related pathways were downregulated in PMM relative to BMPCs (**Fig. 3D**).

**Fig. 3.**
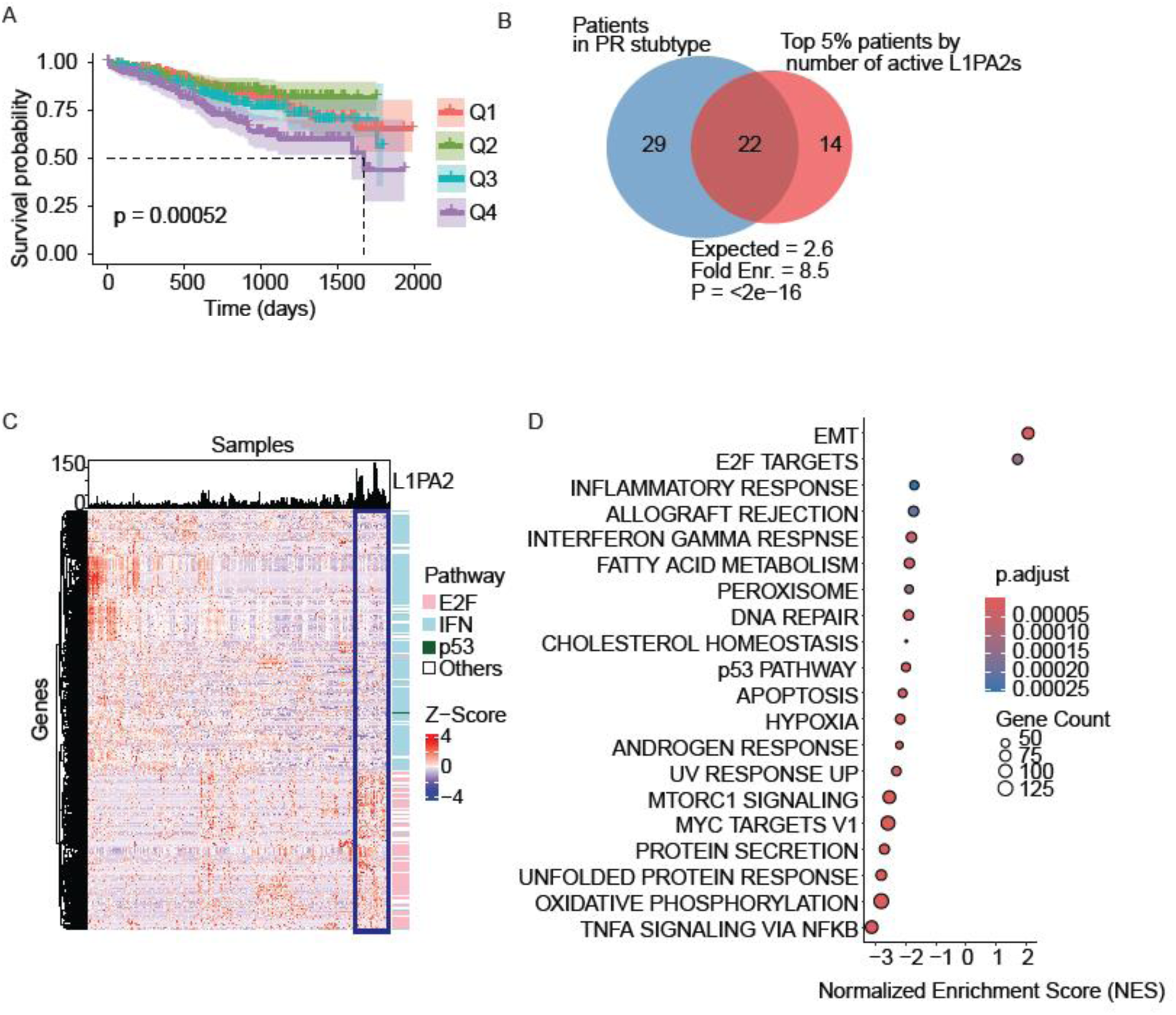
LINE1 activation as a clinical indicator for MM progression. **A**, Kaplan-Meier survival analysis of MM patients stratified by number of active L1s (Q1-Q4). **B**, Overlap between patients in the PR subtype and top 5% patients by number of active L1PAs. Among the top 5% of MM patients (n = 36) ranked by L1PA2 activation, 22 patients overlapped with the PR subtype (n = 51), which is associated with a high proliferation index. **C**, Unsupervised clustering of differentially expressed genes in MM patients based on L1PA2 activation scores, revealing E2F pathway activation and inhibition of IFN and P53 pathways in patients with high L1PA2 activation. **D**, Pathway analysis for PMM samples also revealed inhibition of TNF, P53, and NF-κB pathways in PMM samples with increased LINE1 activation compared to BMPCs.

To investigate the genes related to patient survival, hazard ratios were calculated for each gene in MM samples. A total of 2876 genes were identified as correlated with prognosis (FDR < 0.05), with 43 of these being KZFPs and 5 of these bona fide L1 binding KZFPs: ZNF93, ZNF519, ZNF248, ZNF425 and ZNF33A (**Supplementary Fig. S3B)**. ZNF93 and ZNF519, associated with poor prognosis, show correlated gene expression patterns across MM samples (**Supplementary Fig. S3C)**. ZNF248, ZNF425 and ZNF33A, associated with good prognosis, also have correlated gene expression patterns across MM samples (**Supplementary Fig. S3C).** Unsupervised clustering of MM samples based on just these five KZFPs resulted in a high-risk and low-risk cluster (**Supplementary Fig. S3D, E)**. These prognosis-related KZFPs show a similar trend in PMM samples. For example, PMM4 tends to express more poor-prognosis KZFPs and fewer good-prognosis KZFPs, suggesting that these KZFPs may have prognosis-related functions in both MM and PMM (**Supplementary Fig. S3F)**.

### LINE1 activity is associated with reduced expression of specific KZFPs

To investigate the relationship between KZFP expression and LINE1 activation in the PMM samples, we analyzed the expression of KZFPs that have been shown to bind to L1s (36). These KZFPs displayed strong co-expression patterns, with their expression levels being highly correlated across samples (**Fig. 4A**). Furthermore, they exhibited a general downregulation in PMM as compared to BMPCs (**Fig. 4A**). Although ZNF382, ZNF585A, and ZNF793 are all located in Cluster 6 on chromosome 19, they are not uniformly co-expressed. ZNF382 is downregulated in PMM, whereas ZNF585A and ZNF793 are upregulated and co-expressed with the oncogene ZNF93 (**Supplementary Fig. S4A**), indicating that this co-expression is not determined by genomic proximity. Given the heterogeneity of active L1 numbers across PMM samples, we focused on the third co-expression cluster, which includes KZFPs with more variable expression in PMM compared to other co-expression clusters. Within this group, ZNF425, ZNF141, ZNF382, ZNF680, and ZNF28 showed the highest variability, and their reduced expression was observed in PMM samples with high L1PA2 activation (**Fig. 4B**). The KZFP binding density weighted by MACS score (i.e., −log₁₀(p-value)) showed that ZNF382, ZNF141, and ZNF425 exhibited higher binding density to L1PA2s activated in PMM compared to ZNF680 and ZNF28. (**Supplementary Fig. S4B**). Among them, the expression of ZNF141 and ZNF382 was negatively correlated with the number of activated L1PA2s across PMM samples (**Fig. 4C**). Downregulation of KZFPs was not unique to PMM; similar reductions were also found in other cancer types, particularly those originating from reproductive and digestive tissues, based on TCGA data (**Supplementary Fig. S4C**). Closer examination of the top differentially expressed KZFPs revealed they are consistently repressed across multiple cancer types compared to normal controls (**Supplementary Fig. S4D**). Increased DNA methylation at promoter CpG for ZNF141 and ZNF382 was found to negatively correlate with their expression (**Fig. 4D**). A similar inverse relationship between KZFP expression and LINE1 activation was observed in MM. However, the specific KZFPs involved differed between MM and PMM, with ZNF425 being the only gene shared across both contexts (**Fig. 4E**). In MM, active L1-binding KZFPs are evolutionary older compared to those in PMM (**Fig. 4E**). KZFPs binding to L1s also displayed strong co-expression patterns in MM samples, with their expression levels highly correlated across samples (**Supplementary Fig. S4E**). Finally, we calculated a repression score based on KZFP expression levels for each sample weighted by the number of binding sites at L1PA2 elements, which showed a significant negative correlation with the number of active L1PA2s in both PMM and MM samples (**Supplementary Fig. S4F**).

**Fig. 4.**
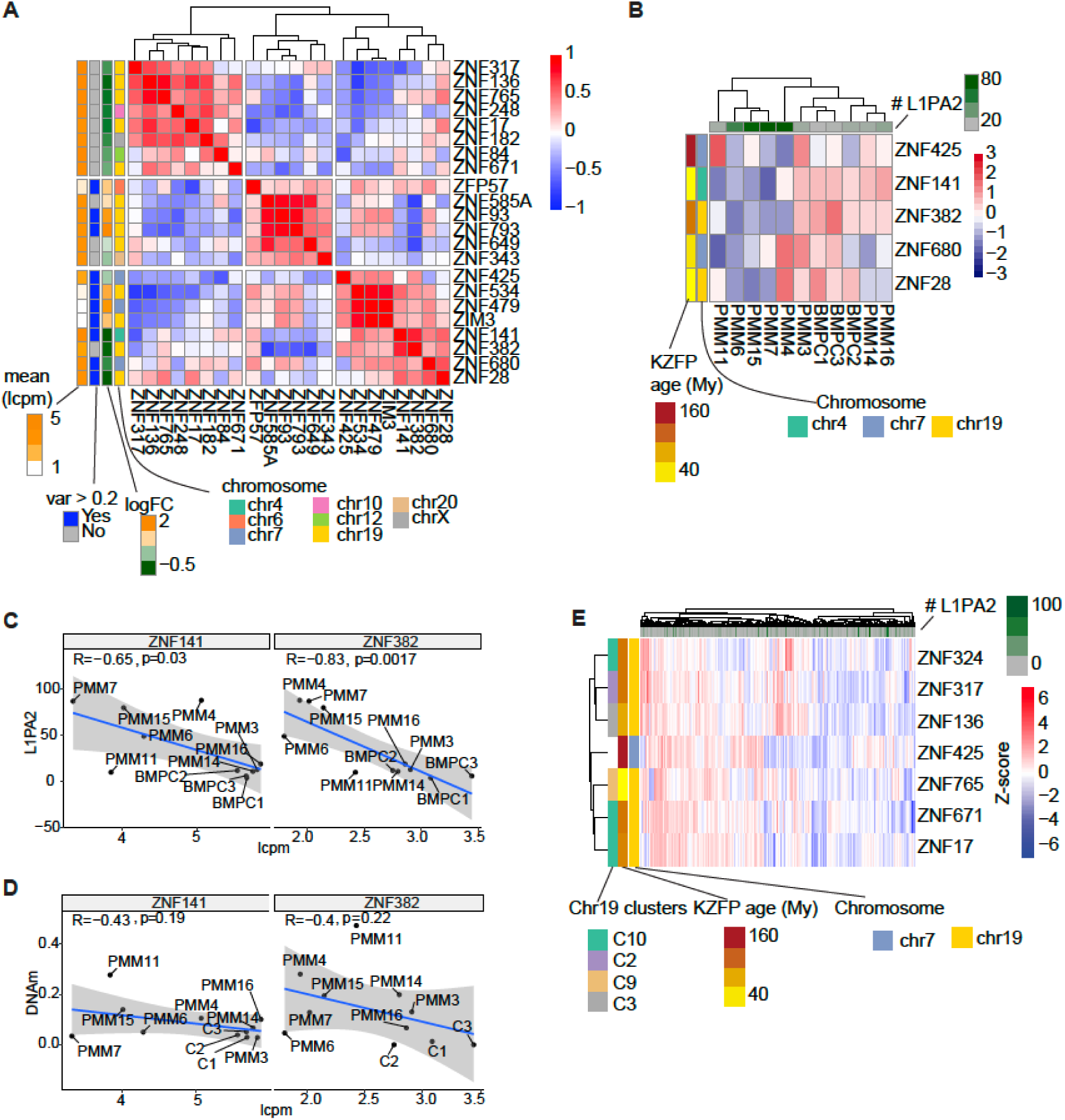
The reduction of ZNF141 and ZNF382 expression is associated to LINE1 activation in PMM. **A**, Correlation of expression levels among KZFPs predicted to bind L1 elements in PMM, filtered for those with more than 10 binding sites at active L1PA2 elements. **B**, Expression levels of KZFPs from Cluster 3 in panel B; samples with lower expression of these genes tend to exhibit higher L1PA2 activation. **C**, Negative correlation between ZNF141 and ZNF382 expression and the number of active L1PA2 transcripts in PMM. **D**, Negative correlation between ZNF141 and ZNF382 expression and DNA methylation levels at their promoter CpG islands (within ±2 kb from TSS) in PMM. **E**, Clustering heatmap of expression for KZFPs negatively correlated with the number of L1s in MM. These KZFPs exhibit an older evolutionary age compared to those in PMM. Six out of seven of these KZFPs were located on chromosome 19, although in different clusters. ZNF425 was found to be common to both MM and PMM.

### L1 activation is impacted by epigenetic changes both in cis and in trans

Given the observed changes in KZFP expression in both PMM and MM, we next aimed to identify potential upstream regulators of KZFP genes. Weighted gene co-expression network analysis was performed using PMM and BMPC samples to identify correlated gene modules. ZNF141 and ZNF382 were clustered in the same gene module and the correlation network showed that genes including IGSF9, DNAJC8P4, BZW1P2, PRL4P2 and NPM1P46 were correlated with both ZNF141 and ZNF382 (**Fig. 5A**). Among these, only IGSF9 is a protein-coding gene, while the others are classified as pseudogenes. IGSF9 expression was positively correlated with ZNF141 and ZNF382 expression and negatively correlated with the number of active L1PA2s (**Fig. 5B**). Since ZNF141 and ZNF382 are expressed at higher levels in embryonic stem cells compared to other cell lines (**Supplementary Fig. S5A**), we re-examined the ChIP-seq data for H1 cells. Analysis of ENCODE ChIP-seq data for H1 cells (37) revealed that several histone-modifying enzymes, including KDM1A, HDAC2, and EZH2, bind to both promoters of ZNF141 and ZNF382 (**Supplementary Fig. S5B**). While the expression levels of KDM1A were not significantly changed in PMM compared to BMPCs, their expression positively correlated with the number of active L1PA2 elements within PMM samples (**Supplementary Fig. S5C**). In addition to KZFP promoter regions, active enhancer regions in H1 cells based on FANTOM5 database were also investigated. We found that DNA methylation levels at enhancers within TE regions were significantly reduced in PMM (**Fig. 5C**). The enhancer whose DNA methylation most strongly correlated with L1PA2 expression in PMM is located within an L1ME5 element (chr7:138243527–138244000) (**Fig. 5D**). This L1ME5 region also overlapped with a KLF6 binding site in the K562 cells (37). Significantly downregulated KLF genes in PMM samples compared to BMPCs included KLF4 and KLF6, two key transcription factors involved in cellular differentiation (**Fig. 5E**). Notably, the same downregulation of KLF4 and KLF6 was also observed in the PR subtype of MM patients, with KLF6 showing a more significant decrease (**Supplementary Fig. S5D**). Although we did not find KLF protein binding sites at full-length L1 regions in the K562 cell line, KLF6 was detected at the 5’UTRs of truncated L1 elements (n=395, **Fig. 5F**). Meanwhile, KLF4 was reported in previous studies to preferentially bind to ERV and SVA repeat families. Interestingly, the transcription factor with the highest binding frequency across the 146 putative functional full-length L1 elements was NRF1 (**Supplementary Fig. S5E**), a gene essential for mitochondrial function. However, NRF1 expression is not differentially expressed in either PMM (**Supplementary Fig. S5F)** or MM samples (**Supplementary Fig. S5G)**. compared to normal control samples.

**Fig. 5.**
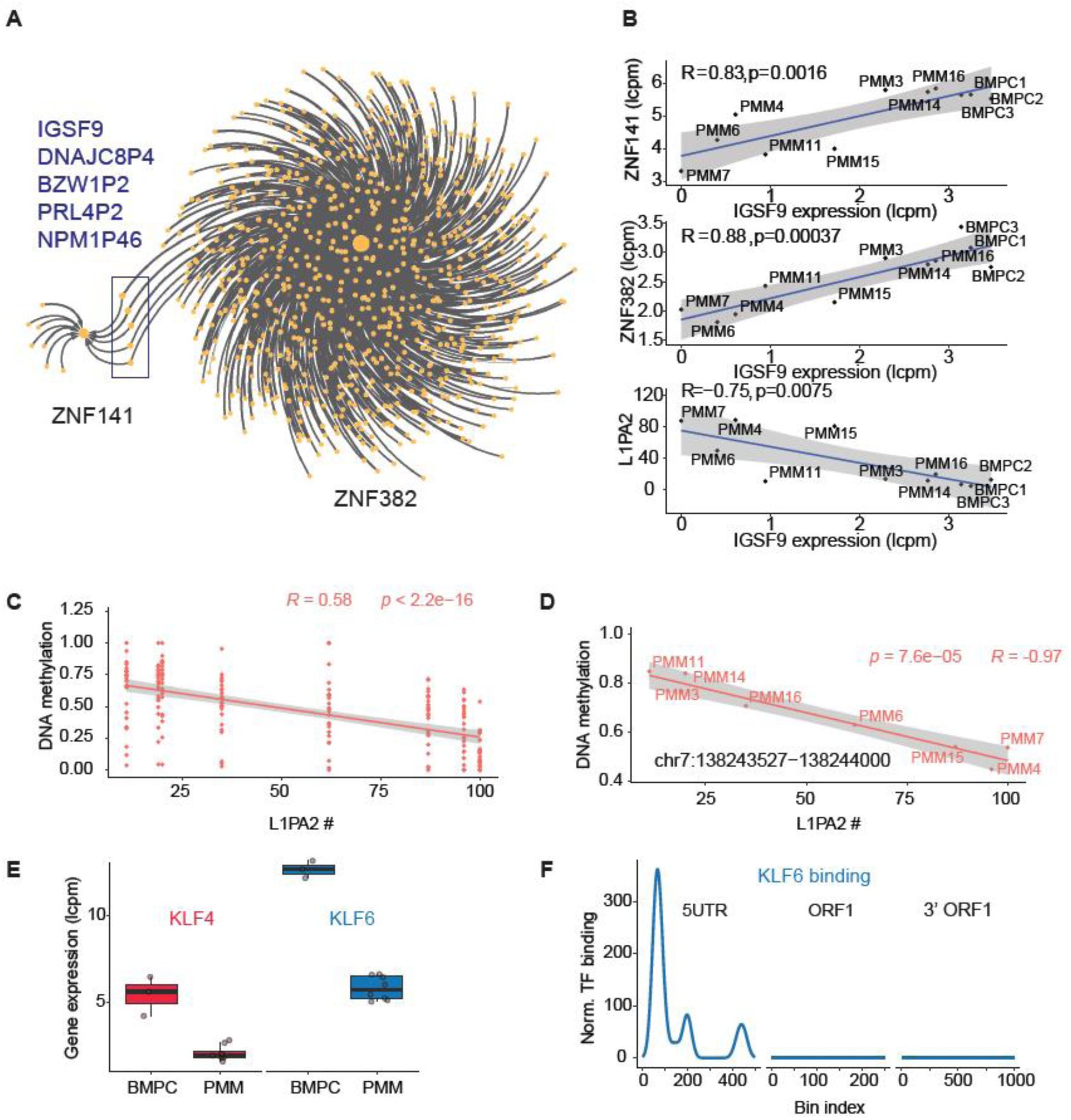
Regulation of L1 activity by potential KZFP regulators and TE enhancers. **A**, Weighted gene co-expression network analysis (WGCNA) revealed that IGSF9, DNAJC8P4, BZW1P2, PRL4P2, and NPM1P46 are commonly positively correlated with both ZNF141 and ZNF382. **B**, The expression of IGSF9 is significantly positively correlated with the expression of ZNF141 and ZNF382, and significantly negatively correlated with the number of active L1PA2 elements. **C**, Enhancers active in H1 cells and located within transposable element (TE) regions have decreased DNA methylation levels in PMM samples with higher L1PA2 activation. **D**, The enhancer with the strongest negative correlation between DNA methylation and L1PA2 activation was located within an L1ME5 element on chromosome 7 (chr7:138243527-138244000). **E**, Expression levels of KLF4 and KLF6 were significantly reduced in PMM compared to BMPC samples (p < 0.05 for *). **F**, KLF6 binding was detected at the 5’UTRs of L1 elements longer than 4500 bp (n = 13,428), most of which lack a complete ORF2. A total of 395 KLF6 binding sites were identified in the K562 cell line.

## Discussion

In this study, we characterized the epigenetic landscape of plasmablastic multiple myeloma (PMM) with a particular focus on DNA methylation changes and the activation of LINE-1 (L1) retrotransposons. Our analysis revealed heterogeneous DNA methylation and aberrant L1 activation across PMM samples, a strong correlation between L1 activity and expression of KRAB zinc finger proteins (KZFPs), and a link between KZFP dysregulation and immune response pathways. These findings highlight the complex interplay between repeat regulation, epigenetic control, and immune signaling in PMM.

L1 activation in PMM likely reflects disruption of normal epigenetic repression mechanisms, including KZFP-mediated DNA methylation and repressive histone modifications (38). When this repression fails, L1s can become transcriptionally active and produce RNA and cDNA intermediates that are recognized by innate immune sensors, such as RIG-I, MDA5, and the cGAS-STING pathway. These events can trigger type I interferon responses, potentially shaping tumor immunogenicity (39). The heterogeneous yet pronounced L1 activation observed in PMM suggests that repeat element dysregulation may contribute to tumor heterogeneity. Intriguingly, cancer cells with pervasive L1 content also have inhibition of interferon and genome pathways, reminiscent of observations in other systems (40,41). Compared to classic multiple myeloma (MM), PMM exhibits a more extensive loss of L1 control, particularly involving IGSF9, which may partially explain its more aggressive clinical course and distinct immune landscape (42). In addition, KLF6 may act as a potential distal activator of KZFPs, with reduced KLF6 expression associated with increased L1 expression in MM and PMM (43).

An unanticipated observation was the differential expression profiles of KZFPs between PMM and MM. Rather than a uniform downregulation of L1PA2 or L1PA3 specific repressors, we observed a complex expression pattern in which some KZFPs are downregulated, while others, such as ZNF93, are upregulated. KZFPs are known to silence TEs by recruiting KAP1/TRIM28 and establishing heterochromatin at TE loci (36). However, emerging evidence suggests that KZFPs may act cooperatively or redundantly, and disruption of this coordination can lead to incomplete or context-dependent TE repression (44). In addition, some KZFPs, including ZNF93, have been implicated in gene regulatory networks and oncogenesis (45), suggesting that KZFP upregulation in PMM may reflect functional adaptation or selection during tumor evolution.

These findings suggest that KZFPs have potential as both biomarkers and therapeutic targets in PMM. Prior studies have shown that TE activity can predict immune checkpoint inhibitor response in certain cancers (46), raising the possibility that KZFP expression profiles could be used to stratify patients based on immune activation or repeat element status. Moreover, pharmacological modulation of repeat repression has shown promise in preclinical cancer models. Drugs that induce L1 expression (e.g., DNA demethylating agents) may convert immune “cold” tumors into “hot” ones by triggering viral mimicry responses (47,48). Conversely, in contexts of chronic inflammation or immune exhaustion, enhancing KZFP-mediated silencing may be beneficial to dampen immune overstimulation (49).

However, several limitations should be noted. Most notably, the number of samples from PMM patients was limited, which restricts the generalizability of our conclusions. Given the rarity of PMM, datasets from larger cohorts will be essential to validate the reproducibility and clinical relevance of these findings.

Future investigations should explore the mechanistic basis by which individual KZFPs influence not only TE silencing, but also immune gene regulation and tumor phenotypes. Recent work has demonstrated that some KZFPs directly regulate non-repetitive genes involved in development and immunity (45,50), suggesting broader functional relevance in cancer biology. Functional studies, such as KZFP perturbation in cell lines or in vivo models will be important to determine causality and therapeutic potential.

In conclusion, our results indicate that KZFP dysregulation plays a critical role in shaping the epigenetic and immune landscape of PMM by modulating L1 activity. These findings position KZFPs as potential biomarkers and therapeutic targets in cancers with altered repeat element regulation.

## Acknowledgments

We thank the patients from City of Hope Medical Center for donating samples and sharing their data for medical research. This work was supported by a pilot award from the Molecular and Cellular Biology Cancer Program of the City of Hope Comprehensive Cancer Center as well as a Steven Gordon & Briskin Family Innovation Grant Program. Research reported in this publication included work performed in the Pathology and Integrative Genomics Shared Resource supported by the National Cancer Institute of the National Institutes of Health under grant number P30CA033572.

## Author Contributions

Q.W. conducted data analyses, performed data visualization, and drafted the manuscript. D.E.S. conceptualized the study, performed data analyses, and revised the manuscript. J.Y.S. provided tissue samples and patient information and contributed to manuscript revision. H.K. generated sequencing libraries. A.L. contributed to data analyses. M.V.B. verified and confirmed KZFP-related analyses.

## References

1. Ribourtout B, Zandecki M. Plasma cell morphology in multiple myeloma and related disorders. Morphologie 2015;99:38–62

2. Nakamura K, Smyth MJ, Martinet L. Cancer immunoediting and immune dysregulation in multiple myeloma. Blood 2020;136:2731–40

3. Tarte K, Zhan F, De Vos J, Klein B, Shaughnessy J, Jr. Gene expression profiling of plasma cells and plasmablasts: toward a better understanding of the late stages of B-cell differentiation. Blood 2003;102:592–600

4. Greipp PR, Raymond NM, Kyle RA, O’Fallon WM. Multiple myeloma: significance of plasmablastic subtype in morphological classification. Blood 1985;65:305–10

5. Kabat M, Patel V, Bhattacharyya P. Plasmablastic multiple myeloma presenting as a pleural mass: a diagnostic challenge. BMJ Case Rep 2024;17

6. Ordonez R, Kulis M, Russinol N, Chapaprieta V, Carrasco-Leon A, Garcia-Torre B, et al. Chromatin activation as a unifying principle underlying pathogenic mechanisms in multiple myeloma. Genome Res 2020;30:1217–27

7. Edwards JR, Yarychkivska O, Boulard M, Bestor TH. DNA methylation and DNA methyltransferases. Epigenetics Chromatin 2017;10:23

8. Lander ES, Linton LM, Birren B, Nusbaum C, Zody MC, Baldwin J, et al. Initial sequencing and analysis of the human genome. Nature 2001;409:860–921

9. de Tribolet-Hardy J, Thorball CW, Forey R, Planet E, Duc J, Coudray A, et al. Genetic features and genomic targets of human KRAB-zinc finger proteins. Genome Res 2023;33:1409–23

10. Babaian A, Mager DL. Endogenous retroviral promoter exaptation in human cancer. Mob DNA 2016;7:24

11. McKerrow W, Wang X, Mendez-Dorantes C, Mita P, Cao S, Grivainis M, et al. LINE-1 expression in cancer correlates with p53 mutation, copy number alteration, and S phase checkpoint. Proc Natl Acad Sci U S A 2022;119:e2115999119

12. Larrayoz M, Garcia-Barchino MJ, Celay J, Etxebeste A, Jimenez M, Perez C, et al. Preclinical models for prediction of immunotherapy outcomes and immune evasion mechanisms in genetically heterogeneous multiple myeloma. Nat Med 2023;29:632–45

13. Solimando AG, Da Via MC, Bolli N, Steinbrunn T. The Route of the Malignant Plasma Cell in Its Survival Niche: Exploring “Multiple Myelomas”. Cancers (Basel) 2022;14

14. Dobin A, Davis CA, Schlesinger F, Drenkow J, Zaleski C, Jha S, et al. STAR: ultrafast universal RNA-seq aligner. Bioinformatics 2013;29:15–21

15. Broad I. Picard Toolkit. 2019

16. Liao Y, Smyth GK, Shi W. featureCounts: an efficient general purpose program for assigning sequence reads to genomic features. Bioinformatics 2014;30:923–30

17. Krueger F, James F, Ewels P, Afyounian E, Weinstein M, Schuster-Boeckler B, et al. FelixKrueger/TrimGalore: v0.6.10 - add default decompression path. 0.6.10: Zenodo; 2023.

18. Krueger F, Andrews SR. Bismark: a flexible aligner and methylation caller for Bisulfite-Seq applications. Bioinformatics 2011;27:1571–2

19. Akalin A, Kormaksson M, Li S, Garrett-Bakelman FE, Figueroa ME, Melnick A, et al. methylKit: a comprehensive R package for the analysis of genome-wide DNA methylation profiles. Genome Biol 2012;13:R87

20. Nishimura D. RepeatMasker. Biotech Software & Internet Report 2000;1:36–9

21. Penzkofer T, Jager M, Figlerowicz M, Badge R, Mundlos S, Robinson PN, et al. L1Base 2: more retrotransposition-active LINE-1s, more mammalian genomes. Nucleic Acids Res 2017;45:D68–D73

22. Chen Y, Chen L, Lun ATL, Baldoni PL, Smyth GK. edgeR v4: powerful differential analysis of sequencing data with expanded functionality and improved support for small counts and larger datasets. Nucleic Acids Res 2025;53:gkaf018

23. Ritchie ME, Phipson B, Wu D, Hu Y, Law CW, Shi W, et al. limma powers differential expression analyses for RNA-sequencing and microarray studies. Nucleic Acids Res 2015;43:e47

24. Robinson MD, Oshlack A. A scaling normalization method for differential expression analysis of RNA-seq data. Genome Biol 2010;11:R25

25. Pertea M, Pertea GM, Antonescu CM, Chang T-C, Mendell JT, Salzberg SL. StringTie enables improved reconstruction of a transcriptome from RNA-seq reads. Nat Biotechnol 2015;33:290–5

26. Patro R, Duggal G, Love MI, Irizarry RA, Kingsford C. Salmon provides fast and bias-aware quantification of transcript expression. Nat Methods 2017;14:417–9

27. Babarinde IA, Ma G, Li Y, Deng B, Luo Z, Liu H, et al. Transposable element sequence fragments incorporated into coding and noncoding transcripts modulate the transcriptome of human pluripotent stem cells. Nucleic Acids Res 2021;49:9132–53

28. Langfelder P, Horvath S. WGCNA: an R package for weighted correlation network analysis. BMC Bioinformatics 2008;9:559

29. Shannon P, Markiel A, Ozier O, Baliga NS, Wang JT, Ramage D, et al. Cytoscape: a software environment for integrated models of biomolecular interaction networks. Genome Res 2003;13:2498–504

30. Song Q, Decato B, Hong EE, Zhou M, Fang F, Qu J, et al. A reference methylome database and analysis pipeline to facilitate integrative and comparative epigenomics. PLoS One 2013;8:e81148

31. Dolgalev I. msigdbr: MSigDB Gene Sets for Multiple Organisms in a Tidy Data Format. 2022

32. Yu G, Wang LG, Han Y, He QY. clusterProfiler: an R package for comparing biological themes among gene clusters. OMICS 2012;16:284–7

33. Berman BP, Weisenberger DJ, Aman JF, Hinoue T, Ramjan Z, Liu Y, et al. Regions of focal DNA hypermethylation and long-range hypomethylation in colorectal cancer coincide with nuclear lamina-associated domains. Nat Genet 2011;44:40–6

34. Yano S, Ishiuchi T, Abe S, Namekawa SH, Huang G, Ogawa Y, et al. Histone H3K36me2 and H3K36me3 form a chromatin platform essential for DNMT3A-dependent DNA methylation in mouse oocytes. Nat Commun 2022;13:4440

35. Skerget S, Penaherrera D, Chari A, Jagannath S, Siegel DS, Vij R, et al. Comprehensive molecular profiling of multiple myeloma identifies refined copy number and expression subtypes. Nat Genet 2024;56:1878–89

36. Imbeault M, Helleboid P-Y, Trono D. KRAB zinc-finger proteins contribute to the evolution of gene regulatory networks. Nature 2017;543:550–4

37. Consortium EP. The ENCODE (ENCyclopedia Of DNA Elements) Project. Science 2004;306:636–40

38. Kannan M, Li J, Fritz SE, Husarek KE, Sanford JC, Sullivan TL, et al. Dynamic silencing of somatic L1 retrotransposon insertions reflects the developmental and cellular contexts of their genomic integration. Mob DNA 2017;8:8

39. You E, Patel BK, Rojas AS, Sun S, Danaher P, Ho NI, et al. LINE-1 ORF1p Mimics Viral Innate Immune Evasion Mechanisms in Pancreatic Ductal Adenocarcinoma. Cancer Discov 2025;15:1063–82

40. Zhao Y, Oreskovic E, Zhang Q, Lu Q, Gilman A, Lin YS, et al. Transposon-triggered innate immune response confers cancer resistance to the blind mole rat. Nat Immunol 2021;22:1219–30

41. Ishak CA, Marhon SA, Tchrakian N, Hodgson A, Loo Yau H, Gonzaga IM, et al. Chronic viral mimicry induction following p53 loss promotes immune evasion. Cancer Discov 2025

42. Moller HE, Preiss BS, Pedersen P, Kristensen IB, Hansen CT, Frederiksen M, et al. Clinicopathological features of plasmablastic multiple myeloma: a population-based cohort. APMIS 2015;123:652–8

43. Pontis J, Planet E, Offner S, Turelli P, Duc J, Coudray A, et al. Hominoid-specific transposable elements and KZFPs facilitate human embryonic genome activation and control transcription in naive human ESCs. Cell Stem Cell 2019;24:724–35.e5

44. Martins F, Rosspopoff O, Carlevaro-Fita J, Forey R, Offner S, Planet E, et al. A cluster of evolutionarily recent KRAB zinc finger proteins protects cancer cells from replicative stress-induced inflammation. Cancer Res 2024;84:808–26

45. Forey R, Pulver C, Raclot C, Rosspopoff O, Offner S, Duc J, et al. Cancer cells subvert the primate-specific KRAB zinc finger protein ZNF93 to control APOBEC3B. bioRxiv 2025:2025.03. 05.641617

46. Zhu X, Fang H, Gladysz K, Barbour JA, Wong JWH. Overexpression of transposable elements is associated with immune evasion and poor outcome in colorectal cancer. Eur J Cancer 2021;157:94–107

47. Chomiak AA, Tiedemann RL, Liu Y, Kong X, Cui Y, Wiseman AK, et al. Select EZH2 inhibitors enhance viral mimicry effects of DNMT inhibition through a mechanism involving NFAT:AP-1 signaling. Sci Adv 2024;10:eadk4423

48. Roulois D, Yau HL, Singhania R, Wang Y, Danesh A, Shen SY, et al. DNA-demethylating agents target colorectal cancer cells by inducing viral mimicry by endogenous transcripts. Cell 2015;162:961–73

49. Ebihara T, Taniuchi I. Exhausted-like Group 2 Innate Lymphoid Cells in Chronic Allergic Inflammation. Trends Immunol 2019;40:1095–104

50. Truby NL, Kim RK, Silva GM, Qu X, Picone JA, Alemu R, et al. A zinc finger transcription factor enables social behaviors while controlling transposable elements and immune response in prefrontal cortex. Transl Psychiatry 2024;14:59

