## Supplementary Methods and Figures for "Methylation variability and LINE-1 activation in multiple myeloma"

**Running title**: Epigenome dynamics and LINE-1 activation in multiple myeloma

***Corresponding Author:**
Dustin E. Schones, Ph.D.
Department of Diabetes Complications and Metabolism
City of Hope
Duarte, CA, 91010


**Supplementary Material**

**Supplementary methods:**

#### Benchmarking for TE expression detection methods

To evaluate methods for transposable element (TE) expression detection, three samples were simulated based on the PMM7 sample, using the same library preparation protocol configured with the CAMPAREE package (1). For each simulated sample, 20 million reads were generated using the BEERS2 package (2). Simulated FASTQ files, containing reads with known alignments to simulated molecules, were then used to benchmark two TE expression detection methods: TEprof3 (3) and the activation score method developed in this study. Transcripts originating from TE regions annotated by RepeatMasker (hg38) served as the ground truth for TE expression. Precision, recall, and F1 scores for LINE-1 transcript detection were calculated at the repeat subfamily level and filtered for only LINE-1 elements with length higher than 5900bp to compare the two methods.

#### Definition and calculation of repression score

The repression score was computed as the weighted sum of selected L1PA2 binding KZFP gene expression levels, with number of KZFP binding regions at L1PA2 sequences serving as the weighting factor. KZFP genes were selected based on co-expression clusters across samples. For each sample, the repression score is calculated with this equation with $i$ means each KZFP:

$$repression\ score\ = \frac{\sum_{i=1}^{n} e_i * b_i}{n}$$

*$e_i$: expression of each KZFP; $b_i$: number of binding regions of each KZFP at active L1PA2 sequences (min-max normalized to a range of 0 to 1).

#### Detection of mobile element insertions (MEI) in PMM

Whole-genome bisulfite sequencing (WGBS) data from PMM and resting B cell samples were first aligned to the CHM13 reference graph genome (https://s3-us-west-2.amazonaws.com/human-pangenomics/pangenomes/freeze/freeze1/minigraph-cactus/hprc-v1.1-mc-chm13/hprc-v1.1-mc-chm13.gfa.gz) using methylGrapher package (4). The insertions (INS) were obtained from pangenome database (https://s3-us-west-2.amazonaws.com/human-pangenomics/pangenomes/freeze/freeze1/minigraph-cactus/hprc-v1.1-mc-chm13/hprc-v1.1-mc-chm13.vcfbub.a100k.wave.vcf.gz), and filtered to retain only those with lengths greater than 50 bp. For each sample, reads aligned to graph nodes covering these INS were extracted from the alignment file. For each INS, read counts and counts per million (CPM) were calculated. All human MEI were downloaded from human genome structural variation consortium version 3 (HGSVC3) from this link: https://ftp.1000genomes.ebi.ac.uk/vol1/ftp/data_collections/HGSVC3/release/Mobile_Elements/1.0/MEI_Callset_T2T-CHM13.ALL.20241211.csv.gz (5). INS overlapped with MEI were selected, and MEIs with CPM higher than 50% quantile of the CPM distribution were selected for valid MEI in PMM samples.

### Analysis of RNA-Seq datasets from the TCGA database

Read counts for all TCGA cancer and normal control RNA-seq datasets were downloaded using the *TCGAbiolinks* R package (6). These read counts were generated from 2-pass STAR alignment of the raw RNA-seq data, as documented on the TCGA portal (https://docs.gdc.cancer.gov/Encyclopedia/pages/STAR_2-Pass_Transcriptome/) and in a previous study (7). The read counts were converted into a list containing sample metadata (for both tumor and normal control samples) and a count matrix, with genes as rows and samples as columns. The data were then normalized using the TMM method (8). Differential expression analysis between tumor and normal groups for each cancer type was performed using the limma-voom pipeline (9,10). KZFP genes were identified from the list of differentially expressed genes, and their corresponding fold changes were used for downstream visualization.

### Hazard Ratio (HR) analysis of MM patient gene expression

To identify prognostically relevant genes, we performed univariate Cox proportional hazards regression on each gene's expression across the MM patient dataset. Clinical data including overall survival time and censoring status were incorporated. HR values, confidence intervals, and *p*-values were computed using the *survival* R package (11). Genes with HR > 1 and FDR < 0.05 were considered associated with poor prognosis.

#### KZFP gene retrieval and clustering

The *biomaRt* R package (12) was used to retrieve all KZFP genes from the Ensembl gene database, as the transcript IDs used in the hg38 genome reference (obtained from the GENCODE database) are Ensembl transcript IDs. Genes were filtered to retain only those containing both C2H2 and KRAB protein domains (Protein IDs: IPR013087 and IPR001909, respectively). Using the *plyranges* R package (13), KZFP gene clusters were defined by applying a minimum intergenic distance (min gap) of 1 Mb, regardless of genomic strand orientation (14).

#### Identification of primate-specific KZFPs

Primate-specific KZFP genes were identified based on prior phylogenetic analysis (15). Genes classified as primate-specific were defined by their presence in the human genome but absence in non-primate mammals, confirmed by comparative genomic alignments and synteny conservation. These genes were extracted from the full KZFP list for focused analysis of species-specific TE repression.

#### Determination of the number of clusters in MM samples based on hazard correlated KZFPs

We applied Gaussian finite mixture modeling using the *mclust* R package (16) to perform model-based unsupervised clustering of the data. The *mclust* package fits a series of Gaussian finite mixture models with varying covariance structures and automatically selects the optimal number of clusters using the Bayesian Information Criterion (BIC). This approach enables the identification of underlying subgroups within the MM samples based on the expression patterns of hazard-related KZFPs.

**Wave score analysis of replication timing**

Replication timing profiles were assessed using wave scores derived from Repli-seq data in HepG2 cells(17). Wave scores represent the signal of early vs. late replication, calculated from fitted curves across the genome. The data were downloaded from the ENCODE project (18) and matched to partially methylated domains to explore the correlation between replication timing, chromatin state, and DNA methylation changes.

**Enrichment analysis of repeat subfamily DNA methylation variation**

To calculate the enrichment score of variable CGIs (defined as variably methylated regions (VMRs) with mean/stdev > 0.2) in repeat families, the observed proportion (numerator of the enrichment score) was defined as the number of variable CGIs within each repeat family divided by the total number of variable CGIs across all TEs. The expected proportion (denominator of the enrichment score) was defined as the number of CGIs (not only VMRs) in each repeat family divided by the total number of CGIs within TEs.

### Publicly available data used in this study

Publicly available RNA-seq datasets were retrieved from the Multiple Myeloma Research Foundation (MMRF) to complement the analysis, samples with complete clinical data and recurrent samples were filtered out. Additional publicly available datasets utilized in this study included ChIP-seq data from the ENCODE database and ChIP-exo data from Imbeault *et al.* (15). Stranded RNA-seq data for BMPC samples were obtained from the GEO database (accession number: GSE114816), and WGBS data for resting B cells were retrieved from GEO (accession number: GSE49627). Wave scores for HepG2 cells were obtained from Thurman *et al.(17)*. KZFP evolutionary age classifications were retrieved from Imbeault *et al.* (15). TEENhancer regions were detected by overlapping with H3K27ac Chip-seq data retrieved from Julien *et al.* (19) with GEO accession number GSE117395 and repeat regions from RepeatMasker (20) and verified using H1 and H9 enhancers in the FANTOM5 database (21).Publicly available datasets used in this study, including TCGA and MMRF-CoMMpass data, were accessed via the GDC Data Portal and the MMRF Research Gateway, respectively, under appropriate data use agreements**.**

### Data visualization

All data visualizations were performed using R (version 4.4.1, released 2024-06-14) or bash scripts. Bar plots and box plots were generated with the *ggplot2* R package (22). Heatmaps were created using either *ComplexHeatmap* (23) or *pheatmap* (24), depending on complexity and annotation needs. Chromosome ideograms were drawn using the *karyoploteR* package (25). Kaplan–Meier survival curves were visualized using the *ggsurvplot* function from the *survminer* R package (26).

**References**


1. Lahens NF, Brooks TG, Sarantopoulou D, Nayak S, Lawrence C, Mrcela A*, et al.* CAMPAREE: a robust and configurable RNA expression simulator. BMC Genomics **2021**;22:692
2. Brooks TG, Lahens NF, Mrcela A, Sarantopoulou D, Nayak S, Naik A*, et al.* BEERS2: RNA-Seq simulation through high fidelity in silico modeling. Brief Bioinform **2024**;25
3. Qu X, Liang Y, McCornack C, Xing X, Schmidt H, Tomlinson C*, et al.* Charting the regulatory landscape of TP53 on transposable elements in cancer. Genome Res **2025**;35:1456-71
4. Zhang W, Macias-Velasco JF, Zhuo X, Belter EA, Jr., Tomlinson C, Garza J*, et al.* methylGrapher: genome-graph-based processing of DNA methylation data from whole genome bisulfite sequencing. Nucleic Acids Res **2025**;53
5. Logsdon GA, Ebert P, Audano PA, Loftus M, Porubsky D, Ebler J*, et al.* Complex genetic variation in nearly complete human genomes. bioRxiv **2024**
6. Colaprico A, Silva TC, Olsen C, Garofano L, Cava C, Garolini D*, et al.* TCGAbiolinks: an R/Bioconductor package for integrative analysis of TCGA data. Nucleic Acids Res **2016**;44:e71
7. Veeneman BA, Shukla S, Dhanasekaran SM, Chinnaiyan AM, Nesvizhskii AI. Two-pass alignment improves novel splice junction quantification. Bioinformatics **2016**;32:43-9
8. Robinson MD, Oshlack A. A scaling normalization method for differential expression analysis of RNA-seq data. Genome Biol **2010**;11:R25
9. Law CW, Chen Y, Shi W, Smyth GK. voom: Precision weights unlock linear model analysis tools for RNA-seq read counts. Genome Biol **2014**;15:R29
10. Ritchie ME, Phipson B, Wu D, Hu Y, Law CW, Shi W*, et al.* limma powers differential expression analyses for RNA-sequencing and microarray studies. Nucleic Acids Res **2015**;43:e47
11. Therneau TM. A Package for Survival Analysis in R 2020.
12. Durinck S, Spellman PT, Birney E, Huber W. Mapping identifiers for the integration of genomic datasets with the R/Bioconductor package biomaRt. Nat Protoc **2009**;4:1184-91
13. Lee S, Cook D, Lawrence M. Plyranges: A grammar of genomic data transformation. Genome Biol **2019**;20:4

14. Martins F, Rosspopoff O, Carlevaro-Fita J, Forey R, Offner S, Planet E*, et al.* A cluster of evolutionarily recent KRAB zinc finger proteins protects cancer cells from replicative stress-induced inflammation. Cancer Res **2024**;84:808-26
15. Imbeault M, Helleboid P-Y, Trono D. KRAB zinc-finger proteins contribute to the evolution of gene regulatory networks. Nature **2017**;543:550-4
16. Scrucca L, Fop M, Murphy TB, Raftery AE. mclust 5: Clustering, Classification and Density Estimation Using Gaussian Finite Mixture Models. R J **2016**;8:289-317
17. Thurman RE, Day N, Noble WS, Stamatoyannopoulos JA. Identification of higher-order functional domains in the human ENCODE regions. Genome Res **2007**;17:917-27
18. Consortium EP. The ENCODE (ENCyclopedia Of DNA Elements) Project. Science **2004**;306:636-40
19. Pontis J, Planet E, Offner S, Turelli P, Duc J, Coudray A*, et al.* Hominoid-specific transposable elements and KZFPs facilitate human embryonic genome activation and control transcription in naive human ESCs. Cell Stem Cell **2019**;24:724-35.e5
20. Nishimura D. RepeatMasker. Biotech Software & Internet Report **2000**;1:36-9
21. Abugessaisa I, Ramilowski JA, Lizio M, Severin J, Hasegawa A, Harshbarger J*, et al.* FANTOM enters 20th year: expansion of transcriptomic atlases and functional annotation of non-coding RNAs. Nucleic Acids Res **2021**;49:D892-D8
22. Ginestet C. ggplot2: Elegant Graphics for Data Analysis. J Roy Stat Soc A **2011**;174:245-
23. Gu Z, Eils R, Schlesner M. Complex heatmaps reveal patterns and correlations in multidimensional genomic data. Bioinformatics **2016**;32:2847-9
24. Kolde R. pheatmap: Pretty Heatmaps. **2025**
25. Gel B, Serra E. karyoploteR: an R/Bioconductor package to plot customizable genomes displaying arbitrary data. Bioinformatics **2017**;33:3088-90
26. Kassambara A, Kosinski M, Biecek P. survminer: Drawing Survival Curves using 'ggplot2'. **2024**

### Supplemental Figures

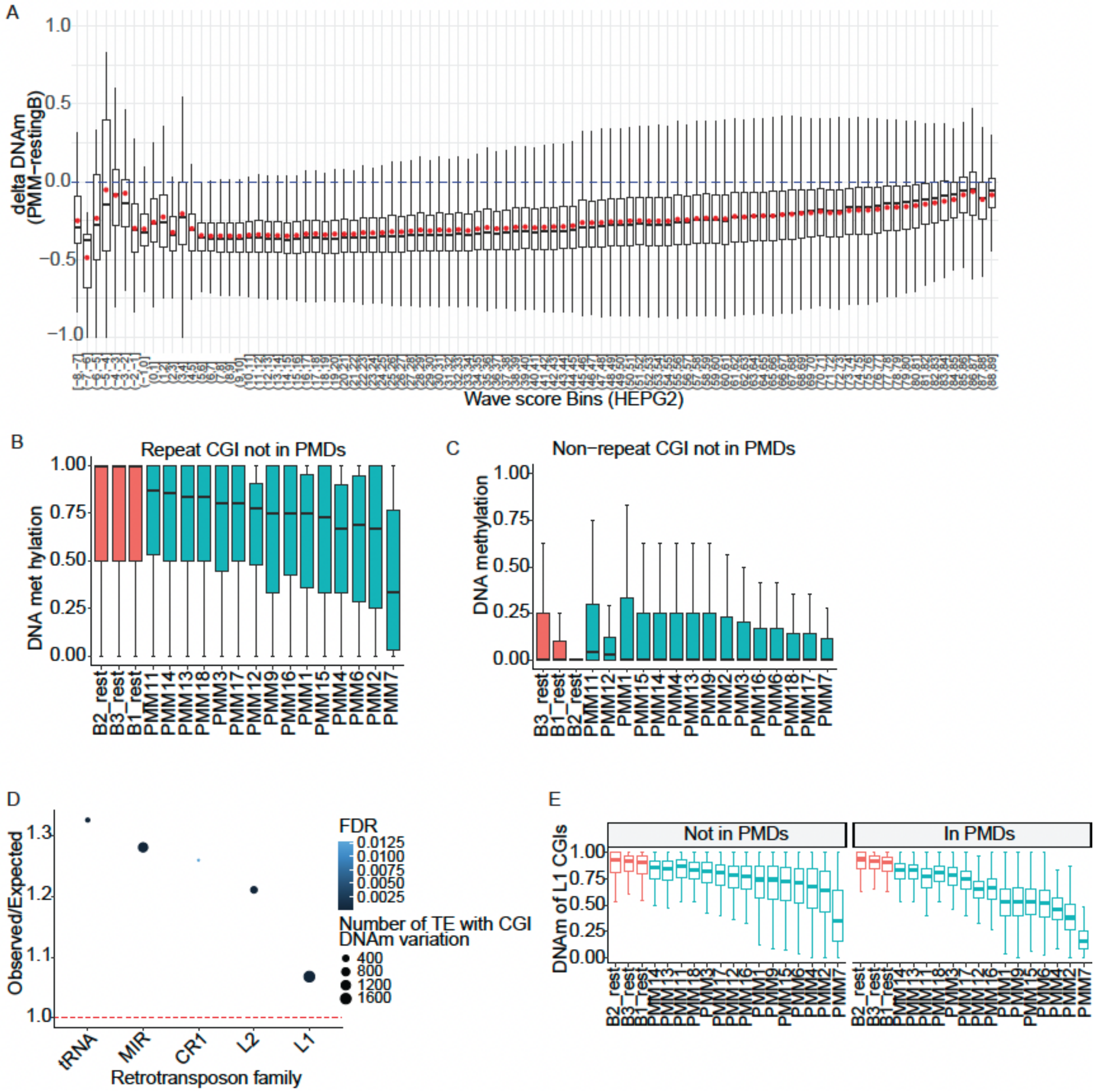


**Supp Fig S1. A**, Boxplots of DNA methylation differences between PMM and resting B cells across wave score bins from HepG2 cells. Late-replicating regions (low wave score) are less methylated in PMM than in resting B cells. Early-replicating regions tend to have similar DNA methylation levels between PMM and resting B cells. **B**, Boxplots of DNA methylation percentage at CGIs in repetitive loci not in PMDs across samples. DNA methylation of these regions is slightly decreased in PMM compared to resting B cells. **C**, Boxplots of DNA methylation percentage at non-repeat CGIs not in PMDs across samples. DNA methylation at these regions is low in both PMM and resting B cells. **D**, Enrichment of repeat families from retrotransposons, including LTRs, LINEs, and SINEs, was performed, and significantly enriched repeat families with CpG island DNA methylation variation (CV > 0.2 and FDR < 0.05) are shown. **E**, Boxplot of DNA methylation of L1 CGIs within and outside of PMDs. DNA methylation decreases more dramatic in PMM for active L1 with CpG islands in PMDs than those not in PMDs.

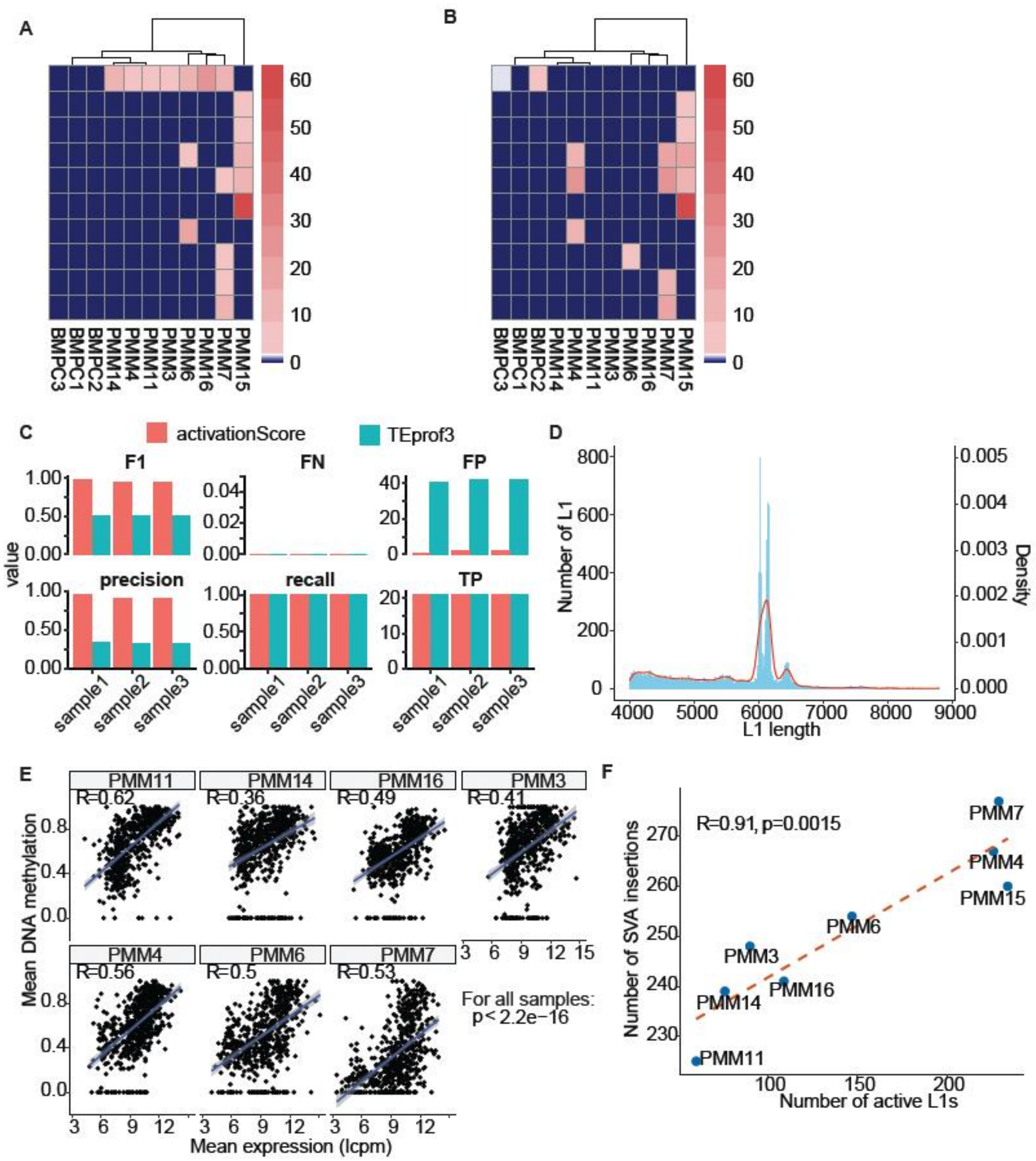


**Supp Fig S2**. Heatmap of showing clustered BMPCs and PMM samples by activation score of active L1s in the (**A**) sense and (**B**) antisense orientation for L1s that are transcribed in both orientations. Only a minority of L1 elements exhibit bidirectional transcription. **C**, Benchmarking the activation score method against TEprof3 and activation score for transposable element (TE) transcript detection indicates that the activation score approach provides higher precision and produces fewer false positives (F1 score is the harmonic mean of precision and recall, calculated as $F1 = 2 \times (precision \times recall) / (precision + recall)$, where $precision = TP/(TP + FP)$ and $recall = TP/(TP + FN)$; TP, FP, and FN are true positives, false positives, and false negatives, respectively.). **D**, Distribution of length for L1s annotated by RepeatMasker (20). **E**, DNA methylation of activated antisense L1s and expression of antisense L1s were positively correlated in PMM samples. **F**, MEIs belonging to the SVA family increase with L1 activation in PMM.

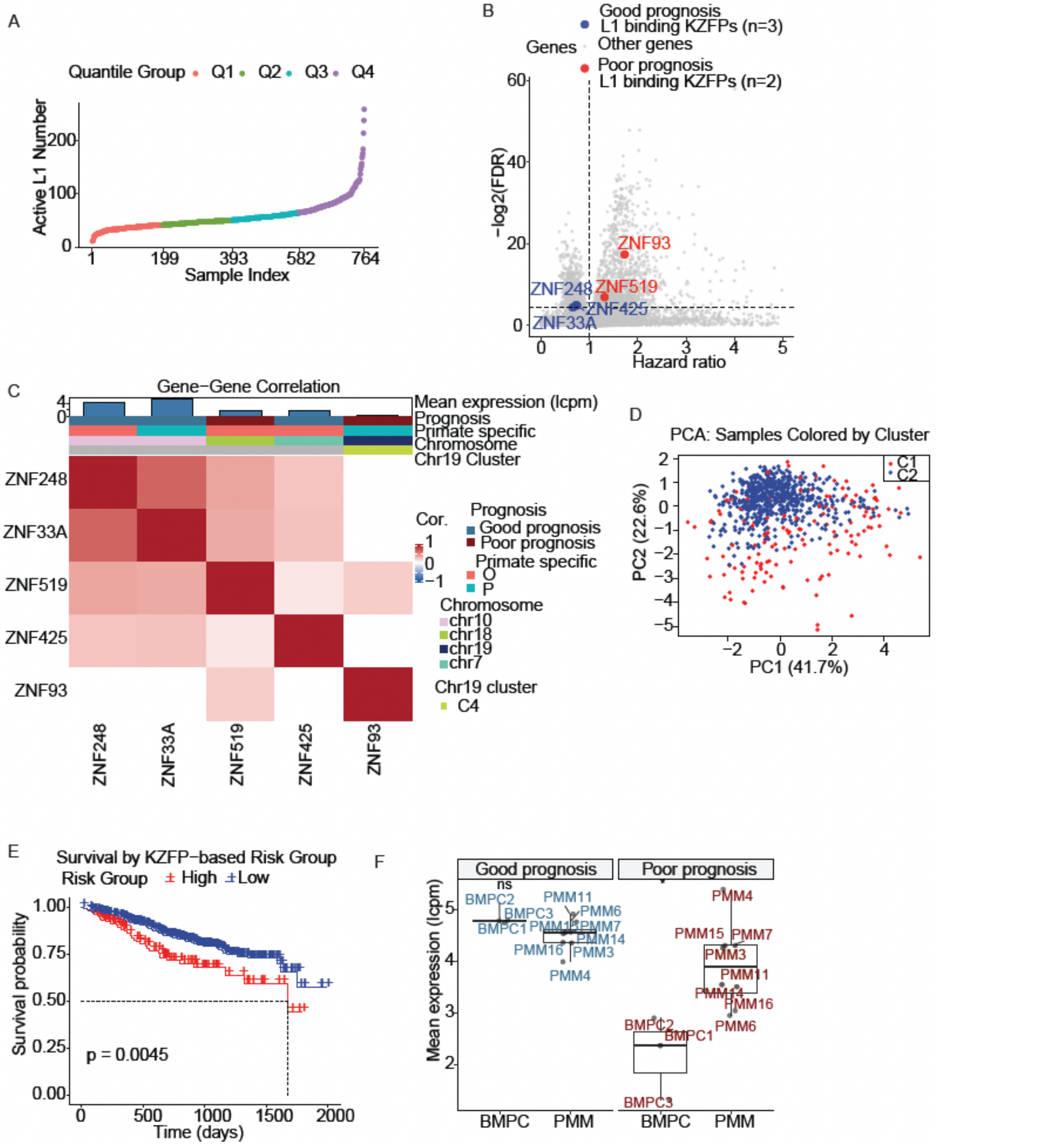


**Supp Fig S3. A**, MM patients were stratified into four quantiles (Q1–Q4) based on the number of active LINE-1 elements, with Q4 representing the group with the highest L1 activation (number of active L1s range from 11 to 259). Shown is a dot plot of active L1 number across the patient cohort. **B**, A volcano plot showed the hazard ratios of all gene expressions in the MM dataset. L1-binding KZFPs significantly associated with prognosis (FDR < 0.05) were labeled in red (n = 2, poor prognosis) and blue (n = 3, good prognosis). **C**, Correlation analysis of the five prognosis-related L1-binding KZFPs revealed that ZNF93 correlated only with ZNF519, and both were associated with poor prognosis. **D**, Unsupervised clustering of MM samples based on the five prognosis-related L1-binding KZFPs divided the samples into two clusters: C1 (red) and C2 (blue). **E**, Survival analysis of samples in C1 and C2 showed that C1 was associated with lower survival probability (red), while C2 was associated with higher survival probability (blue). **F**, Boxplot of gene expression of five prognosis-related KZFPs in PMM samples demonstrates significantly increased expression of the poor prognosis KZFPs (ZNF93 and ZNF519) in PMM compared to BMPC.

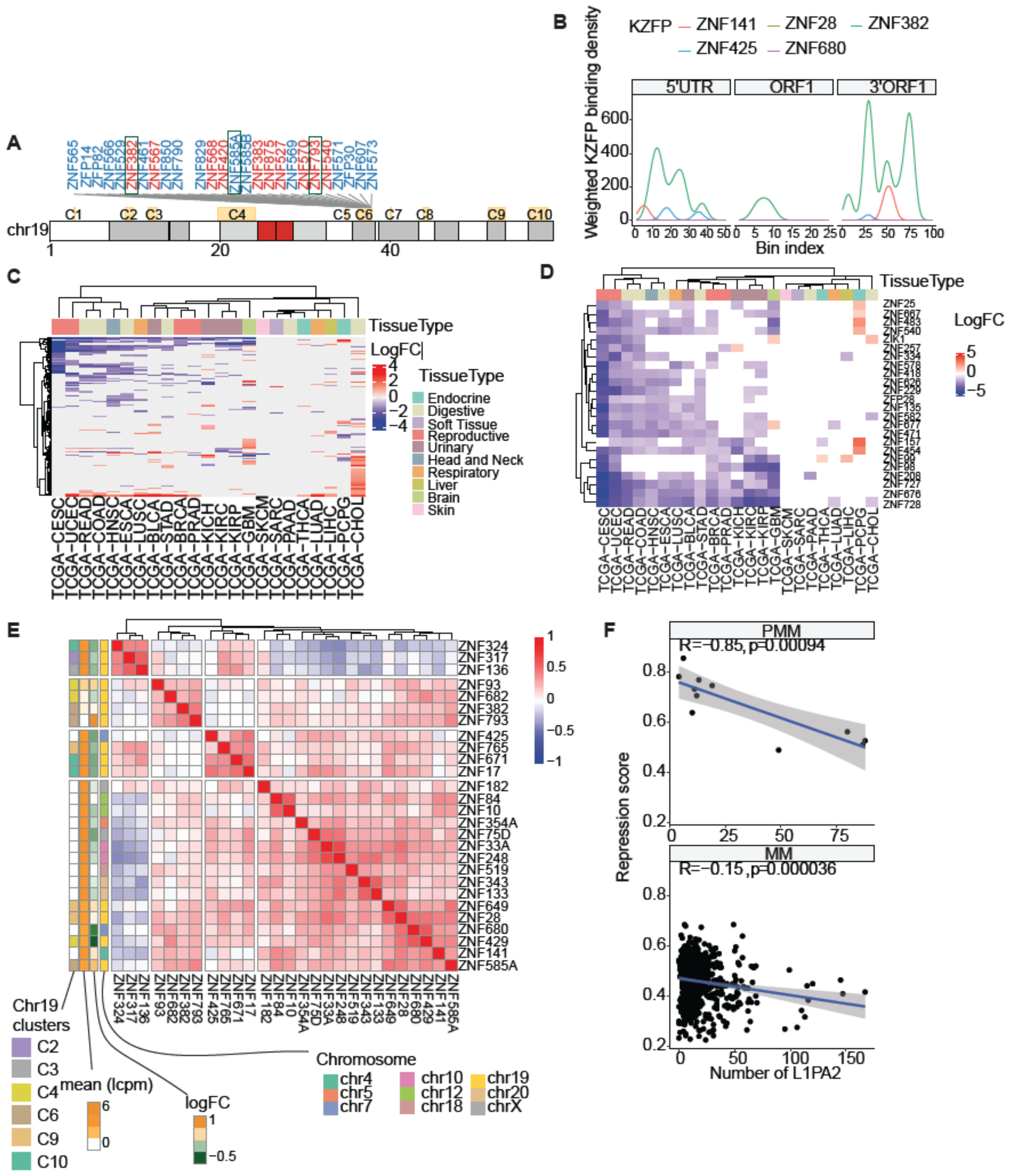

**Supp Fig S4. A**, Gene clusters identified on chromosome 19; genes in Cluster 6 are shown with red indicating genes on the positive strand and blue indicating genes on the negative strand. **B**, Log fold changes of KZFP expression across different cancer types compared to their corresponding normal control tissues using TCGA datasets. **C**, Heatmap illustrating the log fold changes of KZFPs from the first cluster by row in panel B. **D**, Correlation of expression levels among KZFPs predicted to bind L1 elements in MM, filtered for those with more than 10 binding sites at active L1PA2 elements. **E**, Correlation of expression levels among KZFPs predicted to bind L1 elements in MM, filtered for those with more than 10 binding sites at active L1PA2 elements. **F**, Repression score for each sample from PMM and MM patients, based on KZFP expression weighted by the number of L1PA2 binding sites.

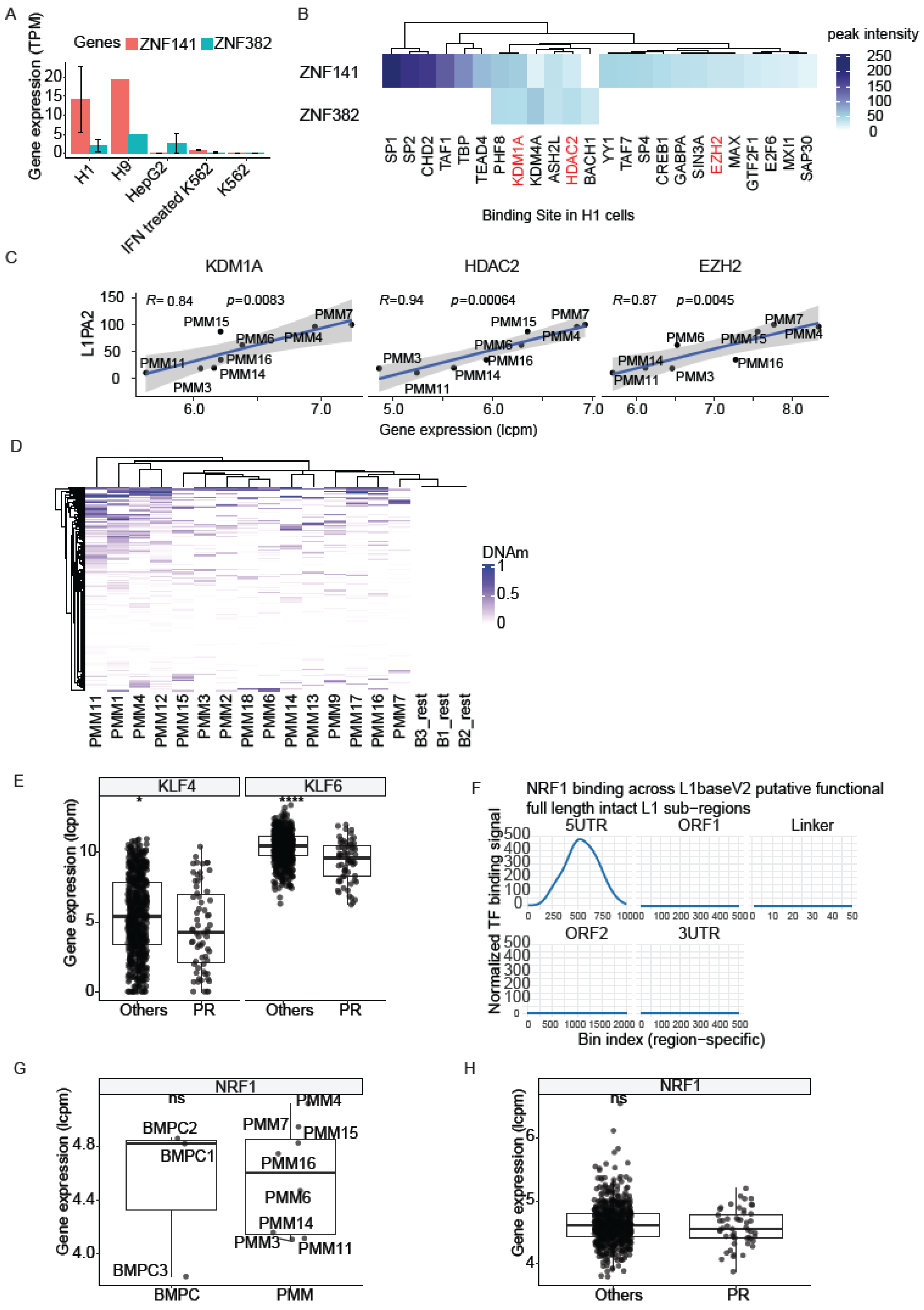
A
Genes ZNF141 ZNF382
Gene expression (TPM)
H1
H9
HepG2
IFN treated K562
K562
B
peak intensity
ZNF141
ZNF382
SP1 SP2 CHD2 TAF1 TBP TEAD4 PHF8 KDM1A KDM4A ASH2L HDAC2 BACH1 YY1 TAF7 SP4 CREB1 GABPA SIN3A EZH2 MAX GTF2F1 E2F6 MXI1 SAP30
Binding Site in H1 cells
C
KDM1A
HDAC2
EZH2
R= 0.84 p=0.0083
R=0.94 p=0.00064
R=0.87 p=0.0045
L1PA2
Gene expression (lcpm)
D
DNAm
PMM11 PMM1 PMM4 PMM12 PMM15 PMM3 PMM2 PMM18 PMM6 PMM14 PMM13 PMM9 PMM17 PMM16 PMM7 B3_rest B1_rest B2_rest
E
KLF4
KLF6
Gene expression (lcpm)
Others
PR
F
NRF1 binding across L1baseV2 putative functional full length intact L1 sub−regions
5UTR
ORF1
Linker
ORF2
3UTR
Normalized TF binding signal
Bin index (region−specific)
G
NRF1
ns
Gene expression (lcpm)
BMPC
PMM
H
NRF1
ns
Gene expression (lcpm)
Others
PR

**Supp Fig S5. A**, Gene expression data from the ENCODE database for H1, H9, HepG2, and K562 cell lines showed that the embryonic stem cell lines H1 and H9 have higher expression of ZNF141 and ZNF382 compared to HepG2 and K562. **B**, Transcription factors (TFs) with significant binding signals (FDR < 0.05) at the promoter regions of ZNF141 and ZNF382, based on ChIP-seq data from H1 embryonic stem cells showed PHF8, KDM1A, KDM4A, ASH2L, HDAC2 and BACH bind to both ZNF141 and ZNF382. **C**, Expression levels of KDM1A, HDAC2 and EZH2 were positively correlated with the number of active L1PA2 elements in PMM samples. **D**, Same as PMM samples, in MM patients of the PR subtype, both KLF4 and KLF6 were downregulated ($p < 0.05$ for *, $p < 0.0001$ for ****). **E**, NRF1 was found to bind to the 5'UTR regions of full-length L1 elements in the K562 cell line, but NRF1 didn't change at gene expression level in both PMM (**F**) and PR subtype of MM (**G**).
